# Empirical decomposition of the explained variation in the variance components form of the mixed model

**DOI:** 10.1101/2019.12.28.890061

**Authors:** Nicholas Schreck

## Abstract

The coefficient of determination is a standard characteristic in linear models. It is widely used to assess the proportion of variation explained, to determine the goodness-of-fit and to compare models with different covariates. However, there has not been an agreement on a similar quantity for the class of linear mixed models yet. We introduce a natural extension of the well-known adjusted coefficient of determination in linear models to the variance components form of the linear mixed model. We propose a novel coefficient of determination which is dimensionless, has an intuitive and simple definition in terms of variance explained, is additive for several random effects and reduces to the adjusted coefficient of determination in the linear model. To this end, we prove a full decomposition of the sum of squares of the independent variable into the explained and residual variance. Based on the restricted maximum likelihood equations, we introduce a novel measure for the explained variation which we allocate specifically to the contribution of the fixed and the random covariates of the model. We illustrate on two genomic datasets that the novel coefficient of determination can be applied as an improved estimator of the heritability of complex traits. Exemplarily, we allocate the explained variation to the five chromosomes of the model plant *Arabidopsis thaliana* and determine the contribution of each chromosome as well as cross-correlations between the chromosomes to the trait heritability.

## 1 Introduction

Genome-wide association studies focus on finding associations between a complex phenotype and marker loci spread on the whole genome, as well as on studying the most common genetic variation in a population (Astle, 2009). Often, testing single nucleotide polymorphisms in separate ordinary regressions has proven to find useful associations and in some cases even to detect quantitative trait loci (The Welcome Trust Consortium, 2007; Newton-Cheh et al., 2009). This approach, however, suffers from low power due to the multiple testing problem (Visscher et al., 2007), from confounding effects of genome-wide loci due to linkage disequilibrium (Segura et al., 2013), and from spurious correlation, i.e. associations that arise not at or not even near a causal locus (Astle, 2009). Therefore, the linear mixed model

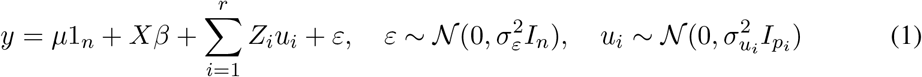

in the traditional variance components form (Searle et al., 1992, §6, §7) is widely applied in genome-wide association studies to fit all markers simultaneously (Kang et al., 2008, 2010; Zhang et al., 2010; Zhou and Stephens, 2012). In model (1), *y* is the vector of observations (e.g. phenotypic values), *μ* ∈ ℝ is an intercept, 1_*n*_ is the column vector of ones of length *n*, *I_n_* denotes the *n*-dimensional identity matrix, and 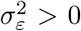 and 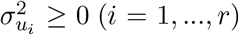. We assume that the design matrix 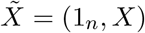 for the fixed effects has full rank (*k* + 1) < *n*. The matrices *Z_i_* (*i* = 1, …, *r*) code covariates with random effects *u_i_*, e.g. genotypes at 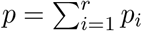 loci.

The heritability of a trait is defined as the proportion of variability in the phenotypic values that can be explained by genomic factors, and is a main quantity of interest in many genomic studies (Falconer and Mackay, 1996). It is well known that the whole genome contributes to the genetic variance for complex traits. It is of central interest to identify genetic variants that explain variation in a phenotype (McCarthy et al., 2008), as well as to attribute the observed variation to specific covariates, in particular to certain regions of the genome such as chromosomes or even single markers (Visscher et al., 2007; Pimentel et al., 2011). In a classical regression context, the heritability equals the coefficient of determination (Falconer and Mackay, 1996), i.e. it measures the strength of the association between the phenotype and the covariates. In more detail, let us first consider the special case

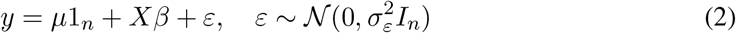

of the linear model (1). The coefficient of determination *R*^2^ = 1 − RSS / TSS is the most popular measure of goodness-of-fit in model (2) (e.g. Kvalseth, 1985), where 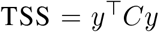 is the centred total sum of squares, 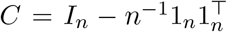 is a centring matrix, 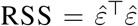 is the residual sum of squares, and 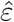 are the estimated residuals, respectively. The centred explained sum of squares is 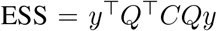 with 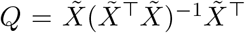. Because of the intercept in model (2) we have TSS = ESS + RSS, which leads to *R*^2^ = ESS / TSS, i.e. the coefficient of determination measures the proportion of total variation that can be explained by the covariates after the least squares fit of model (2), see Kvalseth (1985) or Anderson-Sprecher (1994). We prove in the Supplementary material that 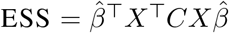, where 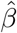 is the least-squares (or maximum likelihood) estimator for *β*. However, the *R*^2^ tends to overestimate the proportion of the explained variation (Kvalseth, 1985). In order to correct for the number of covariates, the adjusted coefficient of determination

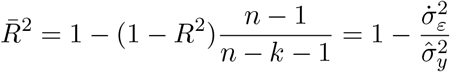

has been introduced. Here, 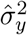 is the empirical variance of *y* and the dot above a variable signifies that restricted-maximum likelihood estimators replace the variance components in the respective quantity. An unbiased estimator for the sample variance 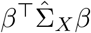 of (*Xβ*)_*i*_ (*i* = 1, …, *n*) is

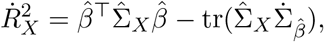

where 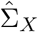 is the empirical covariance matrix of *X* and 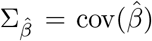. In total, we obtain the decomposition

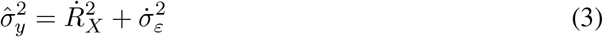

of the centred total sum of squares, and the alternative expression

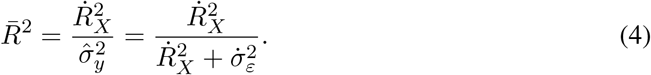

The sample variance 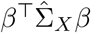 is induced by the empirical variation of the explanatory variables with design *X*, and the estimator 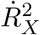 measures the (adjusted) empirical variation explained by the covariates and their empirical pairwise covariances. We derive these results in more detail in the Supplementary material. The 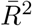 in equation (4) measures the goodness of the model fit, has an intuitive reasonable interpretation in terms of explained empirical variation, is independent of the units of measurement of the model variables, i.e. it is dimensionless, has values in the range of 0 and 1, and equals 1 in case of a perfect fit. These are some of the guidelines for measures of model fit introduced by Kvalseth (1985). Besides its interpretation as a measure of explained variation, the 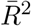 is a wide-spread tool for comparisons of models with a common null model (Anderson-Sprecher, 1994), e.g. a model that contains only the intercept.

The random effects model

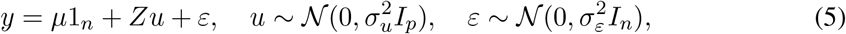

a special case of model (1) with only one random effect and no fixed covariates but the intercept, is used in many genomic applications. Interestingly, it has not been agreed on a unified coefficient of determination for model (5) yet. Typical approaches to define a measure of goodness-of-fit include likelihood-based methods and measures for the total model variance (e.g. Piepho, 2019). In the framework of model (5), the heritability is often defined as

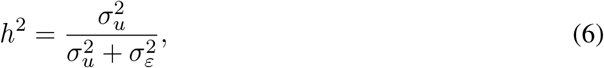

i.e. a ratio of the variance components of the model (e.g. Loh et al., 2015). The estimated variance component 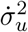 is considered as the explained variation, and the ratio (6) is structurally similar to the 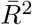 in equation (4). However, the variance components do not completely decompose the phenotypic variation as in equation (3), and it is not possible to fully attribute the total observed variation to the covariates of the model. It has been widely stated that ratios similar to (6) should not be used to estimate the proportion of the variation explained by the genomic variance (Gianola et al., 2009; de los Campos et al., 2015). Recently, Schreck et al. (2019) have introduced the empirical best predictor

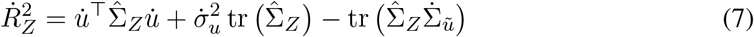

for the empirical genomic variance 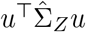. Here, 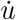 is the empirical best linear unbiased predictor of the random effects *u* and 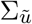 is the covariance matrix of the best linear unbiased predictor 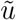. The striking equality

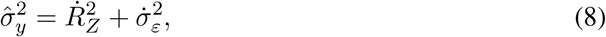

which is similar to equation (3), has been observed by Schreck et al. (2019) in several applications. This suggests a connection between the estimated genomic variance in (7) and the concept of the explained variation in the random effects model similar to model (2).

In the following, we prove a decomposition of the total sum of squares for model (1). This decomposition is based on the wide-spread restricted maximum likelihood estimation of the variance components. We derive a natural and consistent extension of the concept of the coefficient of determination to the variance components form of the linear mixed model, obtaining the equalities (3) and (8) as special cases. More specifically, we decompose the empirical explained variation into separate contributions by the fixed covariates, the random covariates and empirical covariance effects between the fixed and the random covariates. We explicitly attribute the explained variation to the five chromosomes of the model plant *Arabidopsis thaliana* and quantify empirical cross-effects between the chromosomes.

## 2 Decomposition in the Linear Mixed Effects Model

The linear mixed model (1) has pairwise uncorrelated random effects (including the residuals). Let the random vector

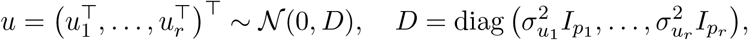

be of dimension *p* and *Z* = (*Z*_1_, …, *Z_r_*) denote an *n × p* design matrix. Model (1) has the variance-covariance structure

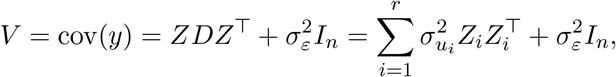

such that 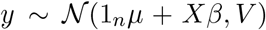. Similar to (Searle et al., 1992, §6), we let 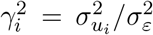 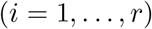 such that 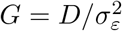, i.e. *G* is *D* but with 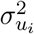 replaced by 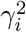. Let 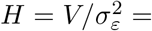 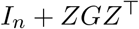 and *H_C_* = *I_n_* + *CZGZ*^T^*C*, which originates from centring the columns of *Z*.

### Assumption 1 (Inverses of matrices).

*We assume that* 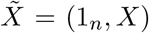 *has full column rank and that the inverses of H*, 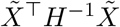, *H*_*C*_ *and* 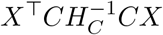 *exist.*

For notational convenience let

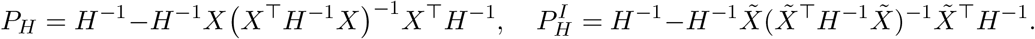

The following lemma states the invariance of the best linear unbiased estimator 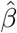 and predictor 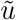 to centring the data *y*, *X*, and *Z*. The invariance also holds for their respective covariance matrices. In particular, the inclusion of a random intercept term in model (1) is not necessary to arrive at this result. In the framework of model (1) this is a striking property, because centring *Z* automatically changes the covariance structure of the model.

### Lemma 2.

*The best linear unbiased estimator of the intercept μ is* 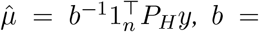 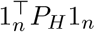, *and the best linear unbiased estimator of the effects β with its covariance matrix is*

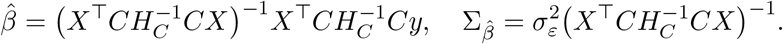

*The best linear unbiased predictor of the random effects u is*

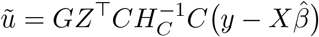

*with covariance matrix*

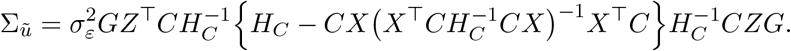

The following lemma introduces unbiased estimators and predictors for the empirical explained variances in model (1).

### Lemma 3.

*Based on model (1) and the quantities introduced in Lemma 2, the estimator*

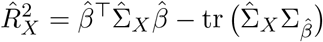

*is unbiased for the sample variance* 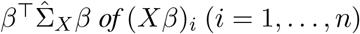, *the predictor*

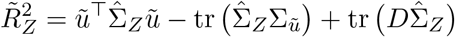

*is unbiased for the sample variance* 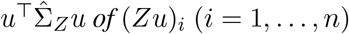, *and*

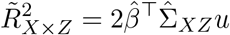

*is an unbiased predictor of the sample covariance* 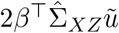 *induced by the possible non-orthogonality of the columns of CX and CZ. The quantities* 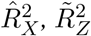, and 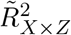 *are invariant to centring the data y, X, and Z*.

The estimation of the variance components is a principal aspect to be considered in the process of fitting model (1). Typically, estimation procedures for the variance components are based on likelihood approaches, where the restricted maximum likelihood approach (Patterson and Thompson, 1971) is preferred in most applications. The restricted maximum likelihood estimates for the variance components are the solution to a system of non-linear equations.

### Lemma 4.

*The set of REML equations (Searle et al., 1992*, §*6)*

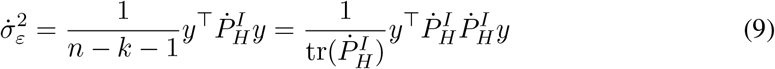

*and*

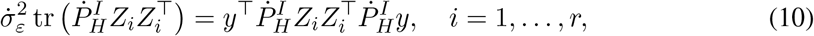

*is invariant to centring y, X and Z. In addition, the equation for the residual variance can be reformulated in the following three ways:*

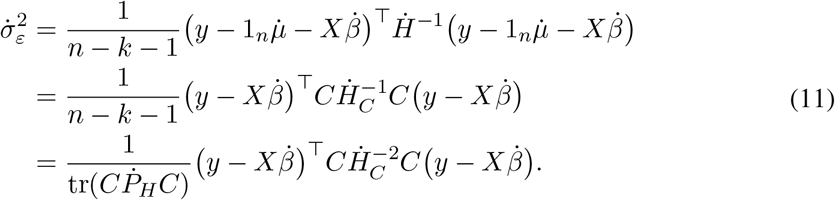

From now on, we assume that we have admissible solutions to the system of equations (9) and (10). We replace the variance components by their estimates and mark the resulting quantities by a dot. In case that the variable already has a superscript, we drop the first superscript for notational convenience. Here, this is only necessary when plugging estimates of the variance components into 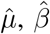 and 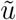. The resulting empirical best linear unbiased estimators 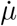 and 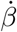 as well as the best linear unbiased predictor 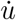 are no longer linear in *y*, and properties such as unbiasedness hold only under additional assumptions (Kackar and Harville, 1984; Jiang, 1999). The following theorem states the empirical decomposition of the centred total sum of squares.

### Theorem 5.

*Given restricted maximum likelihood estimators* 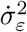 *for the residual variance* 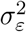 *and* 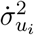 *for the variance components* 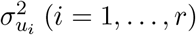 *in model (1), we find*

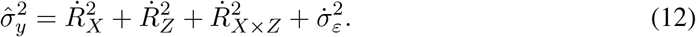

An intuitive reasoning for variables 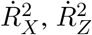, and 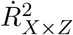 is given by Lemma 3. A proof of this theorem can be found in the Appendix. Similar to 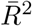 in equation (4), let 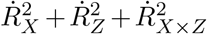 be the empirical explained variation and

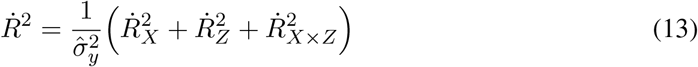

be a coefficient of determination for the linear mixed model (1). The additive decomposition derived in (12) gives

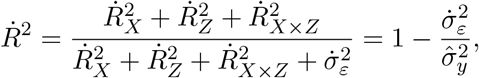

i.e. the possible range of values of 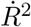 is [0, 1] and it equals 1 if the residual variance equals 0. The quantity 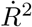 is dimensionless, i.e. it is invariant to scaling and centring the data *y*, *X*, and *Z*. The 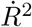 behaves additively for several random effects because

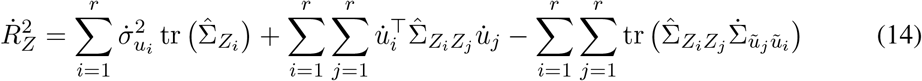

and

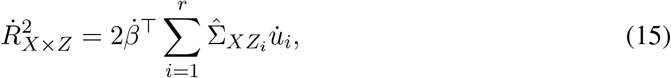

i.e. we can allocate the explained variation to specific contributions of covariates and empirical covariances between covariates. The definition of 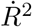 in (13) reduces to 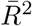 in the linear model (2), i.e. it is a natural extension of the adjusted coefficient of determination from model (2) to model (1). We obtain the variance decomposition (8) in case of the random effects model (5), see also Schreck et al. (2019). The 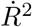 is an analytic function of the estimates returned after a model fit with restricted maximum likelihood so that its computation is straightforward and fast. Maybe even more importantly, due to its explicit nature, an interpretation of the origin and the nature of the explained variation is possible. In particular, the explained variation is caused by the sample variation of the covariates, no matter their specific nature. The contribution of each covariate to the explained variation is clearly visible. Equation (12) implies that we can additively attribute the explained variance within model (1) to the fixed effects, the random effects, and a contribution caused by potential sample correlations between the fixed and random effects co-variables.

## 3 Data Analysis

### 3.1 Datasets

We consider the mice dataset that comes with the R package BGLR (Perez and de los Campos, 2014) and originates from experiments by Valdar et al. (2006a,b). The dataset includes *n* = 1814 mice. We consider the phenotype “Obesity Bodyweight” as the dependent variable. In the framework of model (1), we will use “Obesity Bodylength” as a fixed covariate, and the full set of *p* = 10, 346 biallelic genotyped loci as random effects with a common variance component, i.e. we set *r* = 1.

In addition, we consider a dataset for the model organism “Arabidopsis thaliana” which is publicly available (The 1001 Genomes Consortium, 2016). The dataset has *n* = 1057 lines which were genotyped for *p* = 193, 697 single nucleotide polymorphism markers. We relate the phenotype flowering time at 10°*C* to the fixed effect covariates, in this case the longitude and the latitude at which the flowers were picked. We considered the marker genotypes on the *r* = 5 chromosomes of the organism as random variables with common variance components. The five chromosomes were genotyped for *p*_1_ = 47, 518, *p*_2_ = 25, 550, *p*_3_ = 38, 813, *p*_4_ = 33, 240 and *p*_5_ = 48, 576 markers, respectively.

### 3.2 Settings

We conduct the following analysis with the free software R (R Development Core Team, 2018). We use the package sommer (Covarrubias-Pazaran, 2017) to fit the linear mixed model defined in (1). We use the option restricted maximum likelihood in sommer to estimate the variance components of the model, see also Lemma 4. Besides the estimated variance components, the package also provides the quantities defined in Lemma 2. We use this outcome to calculate the variance decomposition introduced in Theorem 5. For simplicity and comparability we scale the sample variance of the dependent variable *y* to be equal to 100, i.e. all the variables that describe empirical variation are automatically relative to the empirical variation in the phenotypes.

### 3.3 Illustration of the empirical decomposition

In Table 1 we illustrate the empirical variance decomposition of the dependent variable and the attribution of the empirical variance to the components of model (1). For both datasets the empirical variance decomposition holds up to at least the sixth digit which is not illustrated for reasons of clarity. The remaining difference is due to the lack of computational accuracy. Compared to *Arabidopsis thaliana*, the residual variance for the mice dataset is considerably large, i.e. the model for *Arabidopsis thaliana* explains a larger part of the variability in the phenotypes. We are able to attribute most of the explained variance to the random effects covariates. The additive genomic variance, and because of the standardized phenotypes also the heritability, is about 30% for the mice dataset and about 86% for the model plant *Arabidopsis thaliana*. We conclude that the phenotype flowering time is highly heritable, whereas the variation in the bodyweight of the mice can be roughly attributed to the variation of body length, heritage, and unknown (environmental) factors in the relation 1 : 2 : 4. Also, the variance explained by the only fixed effect variable, body length, is about half of the variance explained by the 10, 346 random genomic covariates. This is somewhat intuitive, because the body mass index depends on the body length. In both datasets, a minor, but not negligible, part of the variation in the observations stems from sample covariation between the fixed and random covariates.

**Table 1:** Empirical explained variation; each entry is multiplied by 100; The total sum of squares was scaled to 100. All numbers have been rounded to two digits.

| | $\dot{R}_X^2$ | $\dot{R}_Z^2$ | $\dot{R}_{X \times Z}^2$ | $\dot{\sigma}_\varepsilon^2$ | $\Sigma$ |
| --- | --- | --- | --- | --- | --- |
| Mice | 16.90 | 30.21 | -8.22 | 61.11 | 100.00 |
| *Arabidopsis thaliana* | 1.23 | 85.73 | 5.44 | 7.59 | 100.00 |

### 3.4 Attribution of empirical explained variation

In Table 2 we illustrate the decomposition of 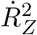 into the contributions of the five chromosomes of *Arabidopsis thaliana* and their cross-effects. No fixed effects but the intercept have been included in the model fit. The formula for the decomposition is given in equation (14), and we attribute

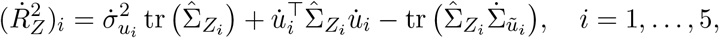

to the single chromosomes. We define the empirical explained covariances analogously by

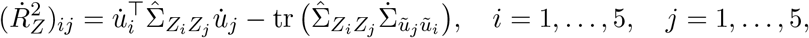

and the correlations by division with the corresponding square roots of the empirical variances.

**Table 2:** Contribution of the five chromosomes of *Arabidopsis thaliana* to the explained variation; first, we depict the 5 × 5 matrix of empirical explained correlations; the last column contains the explained variation by the respective chromosome; All numbers have been rounded to two digits.

| Chromosomes | 1 | 2 | 3 | 4 | 5 | $(\dot{R}_Z^2)_i$ |
| --- | --- | --- | --- | --- | --- | --- |
| 1 | 1.00 | 0.12 | 0.06 | 0.13 | 0.17 | 0.04 |
| 2 | 0.12 | 1.00 | 0.06 | 0.16 | 0.17 | 0.06 |
| 3 | 0.06 | 0.06 | 1.00 | 0.08 | 0.11 | 0.03 |
| 4 | 0.13 | 0.16 | 0.08 | 1.00 | 0.23 | 0.14 |
| 5 | 0.17 | 0.17 | 0.11 | 0.23 | 1.00 | 0.35 |

Chromosome five alone contributes the largest amount to the explained variation, whereas the contributions of the first three chromosomes are comparably small. The empirical explained correlations are the largest between the fifth and the other chromosomes. The magnitude of the correlations indicates the necessity to include them in the analysis. The empirical covariances between the chromosomes contribute about 22% to the observed phenotypic variation.

We infer the direct and indirect contribution of single (fixed and random) components to the explained variation for the mice dataset. We allocate the contribution of each single random effect to the explained variation by setting *r* = 1:

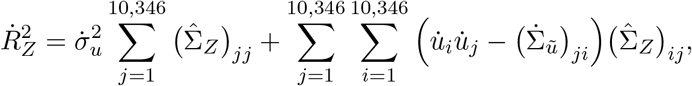

i.e. the explained variation by the random effect covariates can be decomposed into the contribution of the *p* markers. In Figure 1 we illustrate the spread of the explained variation across the genome of the mice. This illustration is similar to the well-known Manhattan plots, i.e. a graph depicting the *p*-values of the single markers in ordinary regressions. However, Figure 1 is an holistic and relative analysis, i.e. the contributions of each marker are always with respect to the other markers, which contrasts the classical single marker analysis in genome-wide association studies. We obtain a map of the genotyped markers and are able to specify regions of lower and larger contributions to the explained variation. This allows for an overview of the genome with respect to the specific trait, and in a next step the further analysis of certain regions of the genome.

**Figure 1:**
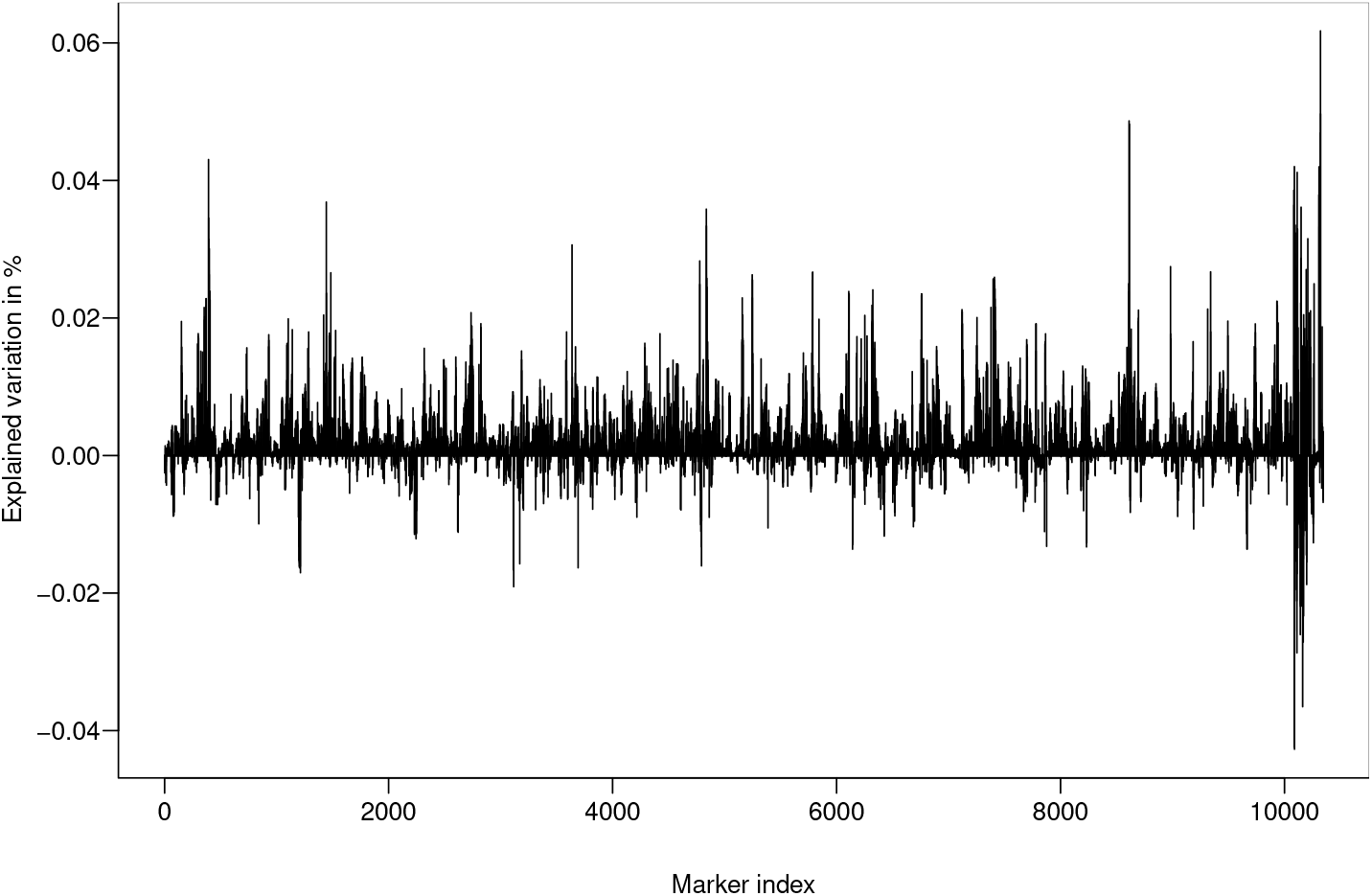
Illustration of the contribution of each marker to the explained variation.

## 4 Discussion

We have proven a novel partitioning of the centred sum of squares in the components form of the linear mixed model, where we have used only estimates that result from the model fit using the restricted maximum likelihood approach. The coefficient of determination 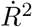 satisfies the most important criteria for appropriate measures of fit, and it is a natural extension of the coefficient of determination from the linear model (2). The decomposition of the total sum of squares derived in Theorem 5 and the associated natural extension of the classical coefficient of determination to the components form of the linear mixed model is in particular important in genomic applications.

We have extended the best prediction approach introduced by Schreck et al. (2019) from the random effects model to the linear mixed effects model. This enables an improved estimation of the additive genomic variance and the heritability for model (1). We have attributed the explained variation, and in particular the genomic variance, to specific components and empirical covariances between covariates of model (1). This decomposition results in graphs that illustrate how the estimated genomic variance is spread across certain regions of the genome. This enables an overview over the relative explained variation by specific (groups of) covariates in a model where these variables are fitted simultaneously. Hence, we have introduced an alternative to the common Manhattan plots, where the variables are tested in separate ordinary regressions.

Variable or model selection is becoming more important with the advent of high-dimensional data. Classically, model comparisons based on (2) are based on the concept of the coefficient of determination, and the model with the larger explained variation is typically chosen. Finding parsimonious models in the framework of model (1) is, however, not straightforward, and there has not been a consensus on how to approach that subject yet (Müller et al., 2013). Approaches include to drop out random covariates with zero variance components (Chen and Brunson, 2003) and a simultaneous selection of fixed and random effects based on a re-parameterized likelihood (Bondell et al., 2010). We investigate the contribution of covariates to the novel coefficient of determination 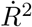. These contributions are not based on the variance components alone, and the sample covariances between different covariates also influence the explained variation. Including these covariances is vital, and can only be neglected if the independent variables are orthogonal to each other. Based on 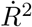 we expect new strategies for variable selection in the framework of model (1).

## Supporting information

Proofs for lemmata

## Acknowledgement

Most sincere thanks are due to Martin Schlather for valuable comments and conversations on this manuscript. I am grateful to Jonas Brehmer and Christopher Dörr for proofreading the manuscript. I was supported by the Deutsche Forschungsgemeinschaft and the Research Training Group 1953.

## Supplementary material

Supplementary Material available online includes the technical details for the empirical variance decomposition in §1, additional technical lemmata for the variance components model and proofs of the lemmata in §2.

## Appendix 1 Proof of the main theorem

*Proof.* (Theorem 5.) First, we make the preliminary considerations that

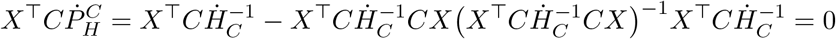

and

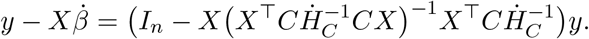

Based on that, we find

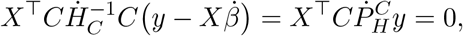

which implies that 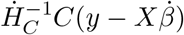 is orthogonal to the column space of *X*. We use this result together with Lemma 2 and the definition of *H_C_* to calculate

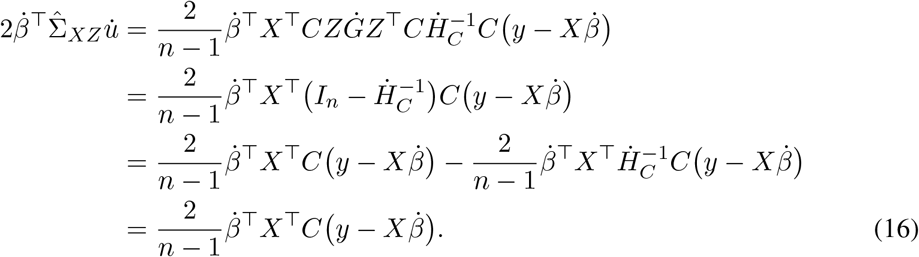

Next, we consider the quadratic form in the empirical best linear unbiased estimators

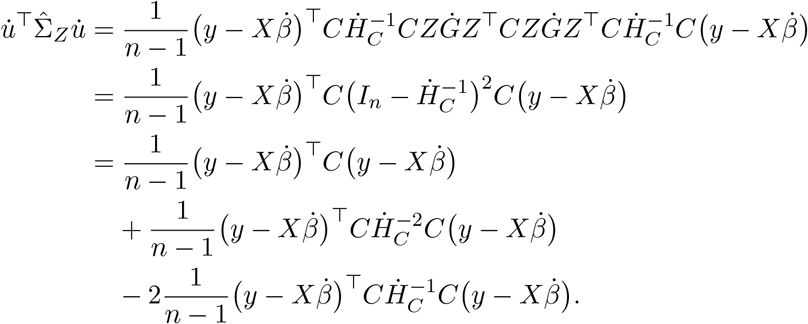

We use equation (11) from Lemma 4 to reformulate

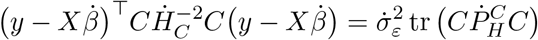

and

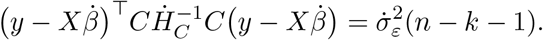

Combining these equations leads to

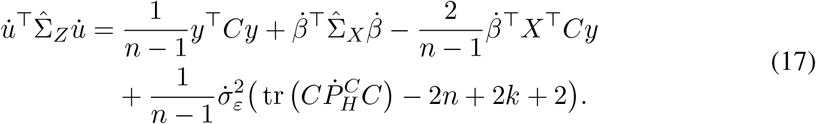

We use Lemma 2 and the cycle property of the trace to calculate

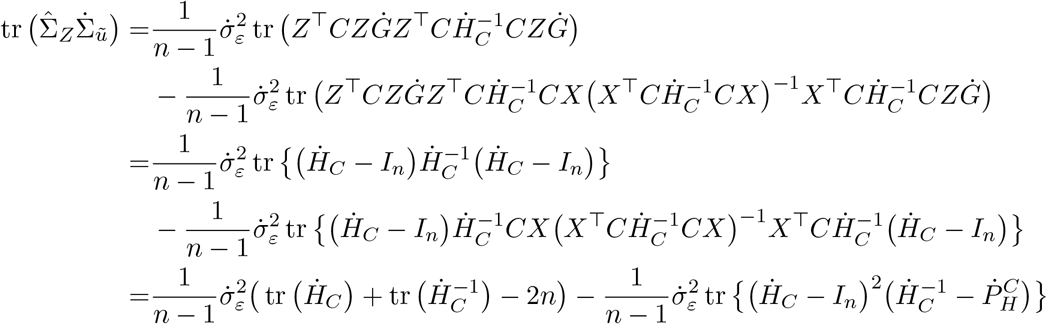

We use

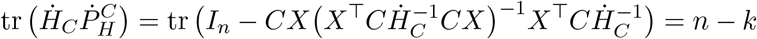

because of the full rank of *k* of *X* to simplify

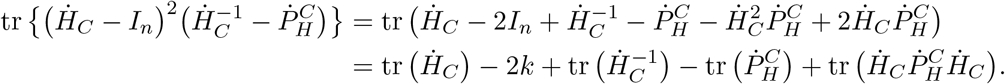

In total, we obtain

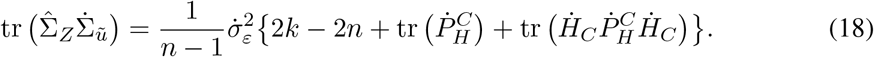

The analogous expression for the fixed effects is

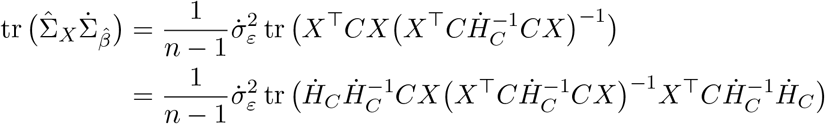

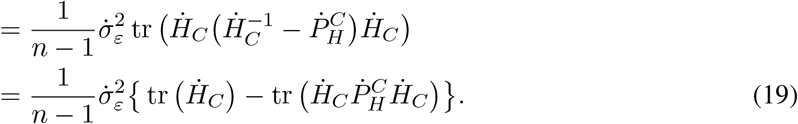

In addition, we consider

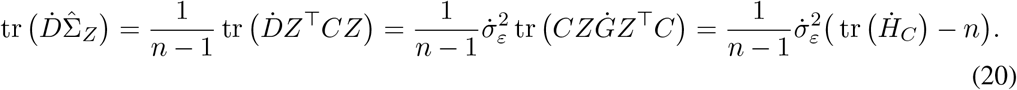

Finally, we use (16), (17), (18) (19), and (20) to calculate the right-hand-side of Lemma 5:

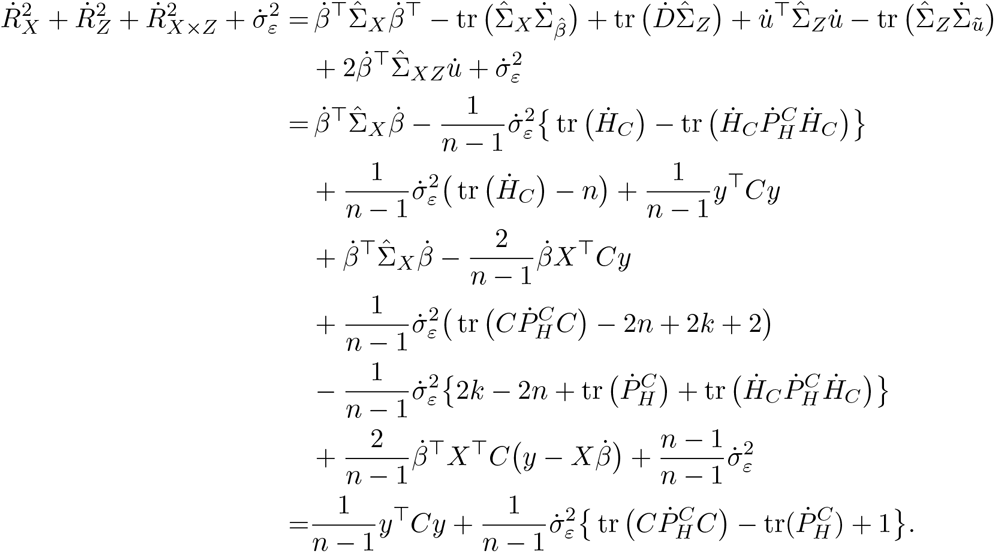

We show in the Supplementary material that 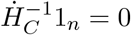. Therefore, we have

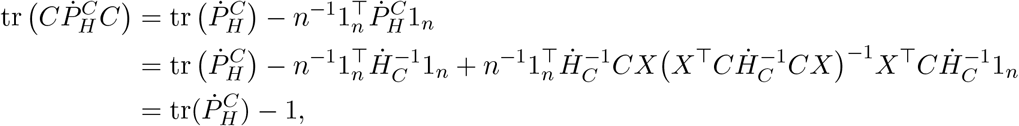

which proves the empirical variance decomposition given in equation (12). The invariance of 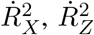 and 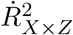 (and with it of 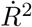) to centering and scaling the columns of *y*, *X*, and *Z* follows by applying Lemmata 2 and 4.

