## Supplementary material for "Empirical decomposition of the explained variation in the variance components form of the mixed model": Proofs for lemmata

**Nicholas Schreck**
University of Mannheim, Mannheim, Germany

December 28, 2019

#### 1 Technical details for the empirical variance decomposition in the linear model

In the linear model

$$y = \mu 1_n + X\beta + \varepsilon, \quad \varepsilon \sim \mathcal{N}(0, \sigma_\varepsilon^2 I_n), \tag{1}$$

let $\delta = (\mu, \beta^\top)^\top$ and $P = I_n - X(X^\top X)^{-1}X^\top$. We assume that the design matrix $\tilde{X} = (1_n, X)$ has full rank $(k+1) < n$.

We have $\tilde{X}^\top(I_n - Q) = 0$ with $Q = \tilde{X}(\tilde{X}^\top\tilde{X})^{-1}\tilde{X}^\top$, and because of the intercept we have $(1_n^\top\hat{\varepsilon}, X^\top\hat{\varepsilon})^\top = \tilde{X}^\top(I_n - Q)y = 0$. Let $\hat{y} = Qy$ be the predicted values. Then, we have the decomposition

$$\begin{aligned}
\text{TSS} &= (y-\bar{y})^\top(y-\bar{y}) = (y-\hat{y})^\top(y-\hat{y}) + (\hat{y}-\bar{y})^\top(\hat{y}-\bar{y}) + 2(y-\hat{y})^\top(\hat{y}-\bar{y}) \\
&= \text{RSS} + \text{ESS} + 2\hat{\varepsilon}^\top\left(\tilde{H} - n^{-1}1_n1_n^\top\right)y \\
&= \text{RSS} + \text{ESS}
\end{aligned}$$

of the centered total sum of squares. It follows directly that

$$\begin{aligned}
y^\top Cy &= \hat{y}^\top C\hat{y} + \hat{\varepsilon}^\top\hat{\varepsilon} \\
&= \hat{\mu}^2 1_n^\top C1_n + \hat{\beta}^\top X^\top CX\hat{\beta} + 2\hat{\mu}1_n^\top CX\hat{\beta} + \hat{\varepsilon}^\top\hat{\varepsilon} \\
&= \hat{\beta}^\top X^\top CX\hat{\beta} + \hat{\varepsilon}^\top\hat{\varepsilon}
\end{aligned}$$

which becomes

$$\hat{\sigma}_y^2 = \hat{\beta}^\top\hat{\Sigma}_X\hat{\beta} + \frac{n-k-1}{n-1}\dot{\sigma}_\varepsilon^2 = \hat{\beta}^\top\hat{\Sigma}_X\hat{\beta} - \frac{k}{n-1}\dot{\sigma}_\varepsilon^2 + \dot{\sigma}_\varepsilon^2. \tag{2}$$

The following lemma states that the best linear unbiased predictor for $\beta$ and its covariance matrix are invariant to centring the data $y$ and $X$.

**Lemma 1.** *In model (1), the best linear unbiased estimator for $\mu$ is*

$$\hat{\mu} = \frac{1}{1_n^\top P 1_n} 1_n^\top Py,$$

*with variance* $\operatorname{var}(\hat{\mu}) = \sigma_\varepsilon^2\big(1_n^\top P 1_n\big)^{-1}$. *The best linear unbiased estimator for the effects* $\beta$ *is*

$$\hat{\beta} = \frac{1}{n-1}\hat{\Sigma}_X^{-1} X^\top C y$$

*with covariance matrix* $\Sigma_{\hat{\beta}} = \operatorname{cov}(\hat{\beta}) = \sigma_\varepsilon^2 (n-1)^{-1}\hat{\Sigma}_X^{-1}$.

This lemma follows as a special case of Lemma 1 from the main paper. We use $\Sigma_{\hat{\beta}}$ given in Lemma 1 and the full rank of $X$ to calculate

$$\operatorname{tr}(\hat{\Sigma}_X \dot{\Sigma}_{\hat{\beta}}) = (n-1)^{-1}\dot{\sigma}_\varepsilon^2 \operatorname{tr}(\hat{\Sigma}_X \hat{\Sigma}_X^{-1}) = k(n-1)^{-1}\dot{\sigma}_\varepsilon^2.$$

We set $\dot{R}_X^2 = \hat{\beta}^\top \hat{\Sigma}_X \hat{\beta} - \operatorname{tr}(\hat{\Sigma}_X \dot{\Sigma}_{\hat{\beta}})$ and obtain the decomposition $\hat{\sigma}_y^2 = \dot{R}_X^2 + \dot{\sigma}_\varepsilon^2$. Lemma 1 states that $\dot{R}_X^2$ is invariant to centring the data $y$ and $X$.

### 2 Technical lemmata for the variance component model

We consider the variance components model

$$y = \mu 1_n + X\beta + \sum_{i=1}^{r} Z_i u_i + \varepsilon, \quad \varepsilon \sim \mathcal{N}(0, \sigma_\varepsilon^2 I_n), \quad u_i \sim \mathcal{N}(0, \sigma_{u_i}^2 I_{p_i}) \tag{3}$$

from the main paper. Here, we prove helpful functional relationships between the characteristics $H$, $H_C = I_n + CZGZ^\top C$, $P_H = H^{-1} - H^{-1}X\big(X^\top H^{-1}X\big)^{-1}X^\top H^{-1}$, $P_H^C$ and $P_H^I = H^{-1} - H^{-1}\tilde{X}(\tilde{X}^\top H^{-1}\tilde{X})^{-1}$ introduced in the main paper. The following lemmata also hold if we replace the variance components by the restricted maximum likelihood estimators, i.e. they also hold for $\dot{H}$, $\dot{H}_C$, $\dot{P}_H$, $\dot{P}_H^C$ and $\dot{P}_H^I$.

**Lemma 2.** *In model (3) it holds that*

$$CH_C^{-1} = H_C^{-1}C = CH_C^{-1}C = H^{-1}\big(I_n - a^{-1}1_n 1_n^\top H^{-1}\big)$$

*with* $a = 1_n^\top H^{-1} 1_n$.

*Proof.* We use the Sherwood-Morrison-Woodbury formula (Henderson and Searle, 1981) to calculate

$$H^{-1} = \big(I_n + ZDZ^\top\big)^{-1} = I_n - Z\big(D^{-1} + Z^\top Z\big)^{-1}Z^\top = I_n - ZA^{-1}Z^\top, \tag{4}$$

where $A = D^{-1} + Z^\top Z$. Analogously, we obtain

$$\begin{aligned} H_C^{-1} &= \big(I_n + CZDZ^\top C\big)^{-1} \\ &= I_n - CZ\big(D^{-1} + Z^\top CZ\big)^{-1}Z^\top C \\ &= I_n - CZ\big(D^{-1} + Z^\top Z - n^{-1}Z^\top 1_n 1_n^\top Z\big)^{-1}Z^\top C \\ &= I_n - CZ\big(A - n^{-1}Z^\top 1_n 1_n^\top Z\big)^{-1}Z^\top C. \end{aligned} \tag{5}$$

We apply the Sherwood-Morrison-Woodbury formula again and obtain

$$\begin{aligned}\big(A - n^{-1} Z^\top 1_n 1_n^\top Z\big)^{-1} &= A^{-1} + A^{-1} Z^\top 1_n \big(n - 1_n^\top Z A^{-1} Z^\top 1_n\big)^{-1} 1_n^\top Z A^{-1} \\ &= A^{-1} + a^{-1} A^{-1} Z^\top 1_n 1_n^\top Z A^{-1},\end{aligned} \tag{6}$$

where

$$a = n - 1_n^\top Z A^{-1} Z^\top 1_n = 1_n^\top \big(I_n - Z A^{-1} Z^\top\big) 1_n = 1_n^\top H^{-1} 1_n. \tag{7}$$

Finally, we use the definition of $C$ and $H^{-1}$ to calculate

$$\begin{aligned}C H_C^{-1} &= H_C^{-1} C = C H_C^{-1} C \\ &\overset{(5)}{=} C - CZ\big(A - n^{-1} Z^\top 1_n 1_n^\top Z\big)^{-1} Z^\top C \\ &\overset{(6)}{=} C - CZA^{-1}Z^\top C - a^{-1} CZA^{-1} Z^\top 1_n 1_n^\top Z A^{-1} Z^\top C \\ &\overset{(7)}{=} H^{-1} - \big\{1 + (n-a)n^{-1} + (n-a)^2 a^{-1} n^{-1}\big\} n^{-1} 1_n 1_n^\top \\ &\quad + \big\{1 + (n-a)a^{-1}\big\} n^{-1} 1_n 1_n^\top Z A^{-1} Z^\top \\ &\quad + \big\{1 + (n-a)a^{-1}\big\} n^{-1} Z A^{-1} Z^\top 1_n 1_n^\top - a^{-1} Z A^{-1} Z^\top 1_n 1_n^\top Z A^{-1} Z^\top \\ &= H^{-1} - a^{-1} 1_n 1_n^\top \big(I_n - Z A^{-1} Z^\top\big) + a^{-1} Z A^{-1} Z^\top 1_n 1_n^\top \big(I_n - Z A^{-1} Z^\top\big) \\ &\overset{(4)}{=} H^{-1} - a^{-1}\big(I_n - Z A^{-1} Z^\top\big) 1_n 1_n^\top H^{-1} \\ &\overset{(4)}{=} H^{-1} - a^{-1} H^{-1} 1_n 1_n^\top H^{-1}.\end{aligned}$$

Factorization proves the lemma. □

**Lemma 3.** *In model (3) it holds that*

$$P_H^I = P_H\big(I_n - b^{-1} 1_n 1_n^\top P_H\big)$$

*with* $b = 1_n^\top P_H 1_n$.

*Proof.* We use the definition of $\tilde{X}$ and have

$$\tilde{X}^\top H^{-1} \tilde{X} = \big(1_n^\top, X^\top\big)^\top H^{-1} (1_n, X) = \begin{pmatrix} a & 1_n^\top H^{-1} X \\ X^\top H^{-1} 1_n & X^\top H^{-1} X \end{pmatrix}.$$

The formula

$$\begin{pmatrix} B_{11} & B_{12} \\ B_{21} & B_{22} \end{pmatrix}^{-1} = \begin{pmatrix} (B_{11} - B_{12} B_{22}^{-1} B_{21})^{-1} & -B_{11}^{-1} B_{12} (B_{22} - B_{21} B_{11}^{-1} B_{12})^{-1} \\ -B_{22}^{-1} B_{21} (B_{11} - B_{12} B_{22}^{-1} B_{21})^{-1} & (B_{22} - B_{21} B_{11}^{-1} B_{12})^{-1} \end{pmatrix}$$

for the inversion of block-matrices for nonsingular $B_{11}$ and $B_{22}$ (Henderson and Searle, 1981) gives

$$\begin{pmatrix} a & 1_n^\top H^{-1} X \\ X^\top H^{-1} 1_n & X^\top H^{-1} X \end{pmatrix}^{-1} = \begin{pmatrix} b^{-1} & -a^{-1} 1_n^\top H^{-1} X B^{-1} \\ -b^{-1} (X^\top H^{-1} X)^{-1} X^\top H^{-1} 1_n & B^{-1} \end{pmatrix}, \tag{8}$$

where

$$b = a - 1_n^\top H^{-1} X \big(X^\top H^{-1} X\big)^{-1} X^\top H^{-1} 1_n = 1_n^\top P_H 1_n \tag{9}$$

and

$$B = X^\top H^{-1}\big(I_n - a^{-1} 1_n 1_n^\top H^{-1}\big) X = X^\top C H_C^{-1} C X \tag{10}$$

because of Lemma 2. The Sherwood-Morrison-Woodbury formula gives

$$B^{-1} = \big(X^\top H^{-1} X\big)^{-1} + b^{-1}\big(X^\top H^{-1} X\big)^{-1} X^\top H^{-1} 1_n 1_n^\top H^{-1} X \big(X^\top H^{-1} X\big)^{-1} \tag{11}$$

and therefore

$$H^{-1} X B^{-1} X^\top H^{-1} = H^{-1} - P_H + b^{-1}\big(H^{-1} - P_H\big) 1_n 1_n^\top \big(H^{-1} - P_H\big). \tag{12}$$

We use the symmetry of $\tilde{X}^\top H^{-1} \tilde{X}$ (and the symmetry of its inverse) and obtain

$$b^{-1}\big(X^\top H^{-1} X\big)^{-1} X^\top H^{-1} 1_n = a^{-1}\big(1_n^\top H^{-1} X B^{-1}\big)^\top \overset{(10)}{=} a^{-1}\big(X^\top C H_C^{-1} C X\big)^{-1} X^\top H^{-1} 1_n. \tag{13}$$

Using (8) and (13) leads to

$$\begin{aligned} H^{-1}\tilde{X}(\tilde{X}^\top H^{-1}\tilde{X})^{-1}\tilde{X}^\top H^{-1} = \\ &= b^{-1} H^{-1} 1_n 1_n^\top H^{-1} - b^{-1} H^{-1} 1_n 1_n^\top H^{-1} X \big(X^\top H^{-1} X\big)^{-1} X^\top H^{-1} \\ &\quad - b^{-1} H^{-1} X \big(X^\top H^{-1} X\big)^{-1} X^\top H^{-1} 1_n 1_n^\top H^{-1} + H^{-1} X B^{-1} X^\top H^{-1} \\ &= b^{-1} P_H 1_n 1_n^\top H^{-1} - b^{-1} H^{-1} 1_n 1_n^\top \big(H^{-1} - P_H\big) + H^{-1} X B^{-1} X^\top H^{-1}. \end{aligned} \tag{14}$$

We use (12) and (14) to derive

$$\begin{aligned} P_H^I &= H^{-1} - H^{-1}\tilde{X}\big(\tilde{X}^\top H^{-1}\tilde{X}\big)^{-1}\tilde{X}^\top H^{-1} \\ &= H^{-1} - b^{-1} P_H 1_n 1_n^\top H^{-1} + b^{-1} H^{-1} 1_n 1_n^\top \big(H^{-1} - P_H\big) - H^{-1} X B^{-1} X^\top H^{-1} \\ &= P_H - b^{-1} P_H 1_n 1_n^\top P_H \\ &= P_H\big(I_n - b^{-1} 1_n 1_n^\top P_H\big) = \big(I_n - b^{-1} P_H 1_n 1_n^\top\big) P_H, \end{aligned}$$

which proves the lemma. □

Combining Lemma 3 and the following lemma results in a structurally similar result for $P_H^C = H_C^{-1} - H_C^{-1} C X \big(X^\top C H_C^{-1} C X\big)^{-1} X^\top C H_C^{-1}$ as we obtain from Lemma 2 for $H_C$.

**Lemma 4.** *In model (3) it holds that*

$$C P_H^C C = C P_H^C = P_H^C C = P_H^I.$$

*Proof.* From the definition of $P_H^C$ and Lemma 2 it follows directly that

$$CP_H^C C = CP_H^C = P_H^C C.$$

Next, we use Lemma 2 and (12) to obtain

$$\begin{aligned}
CP_H^C C =& CH_C^{-1}C - CH_C^{-1}CX\big(X^\top CH_C^{-1}CX\big)^{-1}X^\top CH_C^{-1}C \\
=& H^{-1} - a^{-1}H^{-1}1_n1_n^\top H^{-1} \\
& - \big(I_n - a^{-1}H^{-1}1_n1_n^\top\big)H^{-1}XB^{-1}X^\top H^{-1}\big(I_n - a^{-1}1_n1_n^\top H^{-1}\big) \\
=& H^{-1} - a^{-1}H^{-1}1_n1_n^\top H^{-1} - \big(I_n - a^{-1}H^{-1}1_n1_n^\top\big)\big\{H^{-1} - P_H \\
& + b^{-1}(H^{-1} - P_H)1_n1_n^\top(H^{-1} - P_H)\big\}\big(I_n - a^{-1}1_n1_n^\top H^{-1}\big).
\end{aligned}$$

Lemma 3 and definitions (7) and (9) lead to $CP_H^C C = P_H^I$. □

### 3 Proof of lemmata from the main paper

*Proof.* (Lemma 1). The best linear unbiased estimator $(\hat\mu, \hat\beta^\top)^\top = (\tilde X^\top H^{-1}\tilde X)^{-1}\tilde X^\top H^{-1}y$ for $(\mu, \beta^\top)^\top$ in model (3) can be derived using either maximum-likelihood derivatives or the mixed-model equations (e.g. Searle et al., 1992, §7). We use (8) and (10) to obtain

$$\begin{pmatrix}\hat\mu \\ \hat\beta\end{pmatrix} = \begin{pmatrix} b^{-1}1_n^\top H^{-1}y - a^{-1}1_nH^{-1}XB^{-1}X^\top H^{-1}y \\ -b^{-1}\big(X^\top H^{-1}X\big)^{-1}X^\top H^{-1}1_n1_n^\top H^{-1}y + B^{-1}X^\top H^{-1}y\end{pmatrix}.$$

Consequently, using (7), (9) and (10), the best linear unbiased estimator for the intercept equals

$$\begin{aligned}
\hat\mu &= b^{-1}\big[1_n^\top H^{-1} - ba^{-1}1_n^\top\big\{H^{-1} - P_H + b^{-1}(H^{-1} - P_H)1_n1_n^\top(H^{-1} - P_H)\big\}\big]y \\
&= b^{-1}\big[1_n^\top H^{-1} - ba^{-1}\big\{1_n^\top H^{-1} - 1_n^\top P_H + (a-b)b^{-1}1_n^\top(H^{-1} - P_H)\big\}\big]y \\
&= b^{-1}1_n^\top P_H y.
\end{aligned}$$

Using (10) and (13), the best linear unbiased estimator for the fixed effects equals

$$\begin{aligned}
\hat\beta &= \big(X^\top CH_C^{-1}CX\big)^{-1}X^\top\big(-a^{-1}H^{-1}1_n1_n^\top H^{-1} + H^{-1}\big)y \\
&= \big(X^\top CH_C^{-1}CX\big)^{-1}X^\top CH_C^{-1}Cy, \qquad (15)
\end{aligned}$$

where the last equality follows from Lemma 2. Alternatively, using (11), we write

$$\begin{aligned}
\hat\beta =& -b^{-1}\big(X^\top H^{-1}X\big)^{-1}X^\top H^{-1}1_n1_n^\top H^{-1}y + \big(X^\top H^{-1}X\big)^{-1}X^\top H^{-1}y \\
& + b^{-1}\big(X^\top H^{-1}X\big)^{-1}X^\top H^{-1}1_n1_n^\top H^{-1}X\big(X^\top H^{-1}X\big)^{-1}X^\top H^{-1}y \\
=& -b^{-1}\big(X^\top H^{-1}X\big)^{-1}X^\top H^{-1}1_n1_n^\top\big(H^{-1} - H^{-1}X\big(X^\top H^{-1}X\big)^{-1}X^\top H^{-1}\big)y \\
& + \big(X^\top H^{-1}X\big)^{-1}X^\top H^{-1}y \\
=& \big(X^\top H^{-1}X\big)^{-1}X^\top H^{-1}\big(y - b^{-1}1_n1_n^\top P_H y\big)
\end{aligned}$$

$$= \big(X^\top H^{-1} X\big)^{-1} X^\top H^{-1}\big(y - 1_n\hat\mu\big).$$

This leads to

$$\begin{aligned} X\hat\beta &= HH^{-1}X\big(X^\top H^{-1}X\big)^{-1}X^\top H^{-1}\big(y-1_n\hat\mu\big) \\ &= H\big(H^{-1} - P_H\big)\big(y-1_n\hat\mu\big) \\ &= \big(I_n - HP_H\big)\big(y-1_n\hat\mu\big). \end{aligned} \tag{16}$$

The covariance matrix of $\hat\mu$ and $\hat\beta$ equals

$$\begin{aligned} \operatorname{cov}\big\{\big(\hat\mu, \hat\beta^\top\big)^\top\big\} &= \big(\tilde X^\top V^{-1}\tilde X\big)^{-1} \\ &= \sigma_\varepsilon^2 \begin{pmatrix} a & 1_n^\top H^{-1}X \\ X^\top H^{-1}1_n & X^\top H^{-1}X \end{pmatrix} \\ &\overset{(8)}{=} \sigma_\varepsilon^2 \begin{pmatrix} b^{-1} & -a^{-1}1_n^\top H^{-1}X\big(X^\top C H_C^{-1} C X\big)^{-1} \\ -a^{-1}\big(X^\top C H_C^{-1} C X\big)^{-1}X^\top H^{-1}1_n & \big(X^\top C H_C^{-1} C X\big)^{-1} \end{pmatrix}. \end{aligned}$$

The best linear unbiased predictor of $u$ (e.g. Searle et al., 1992, §7) equals $\tilde u = GZ^\top H^{-1}(y - 1_n\hat\mu - X\hat\beta)$. We use (7), (9) and (16) to write

$$\begin{aligned} a^{-1}1_n1_n^\top H^{-1}\big(y - X\hat\beta\big) =& a^{-1}1_n1_n^\top H^{-1}\{y - \big(I_n - HP_H\big)\big(y-1_n\hat\mu\big)\} \\ =& a^{-1}1_n1_n^\top H^{-1}1_n\hat\mu + a^{-1}1_n1_n^\top H^{-1}HP_H y \\ & - a^{-1}1_n1_n^\top H^{-1}HP_H 1_n\hat\mu \\ =& 1_n\hat\mu + a^{-1}1_n1_n^\top P_H y - ba^{-1}1_n\hat\mu \\ =& 1_n\hat\mu. \end{aligned}$$

Together with Lemma 2 this yields

$$\begin{aligned} H^{-1}\big(y - 1_n\hat\mu - X\hat\beta\big) &= H^{-1}\big(y - a^{-1}1_n1_n^\top H^{-1}\big(y - X\hat\beta\big) - X\hat\beta\big) \\ &= H^{-1}\big(I_n - a^{-1}1_n1_n^\top H^{-1}\hat\beta\big)\big(y - X\hat\beta\big) \\ &= CH_C^{-1}C\big(y - X\hat\beta\big), \end{aligned} \tag{17}$$

and therefore

$$\tilde u = GZ^\top H^{-1}\big(y - 1_n\hat\mu - X\hat\beta\big) = GZ^\top CH_C^{-1}C\big(y - X\hat\beta\big). \tag{18}$$

Clearly, $\tilde u$ is invariant to centring $y, X$ and $Z$. Since by (15)

$$\begin{aligned} \operatorname{cov}\big(y - X\hat\beta\big) =& \sigma_\varepsilon^2 H + \sigma_\varepsilon^2 X\big(X^\top CH_C^{-1}CX\big)^{-1}X^\top \\ & - 2\sigma_\varepsilon^2 HCH_C^{-1}CX\big(X^\top CH_C^{-1}CX\big)^{-1}X^\top, \end{aligned}$$

and by definition of $H_C$

$$CHCH_C^{-1}C = CH_CH_C^{-1}C = C, \tag{19}$$

we calculate the covariance matrix $\Sigma_{\tilde{u}} = \mathrm{cov}(\tilde{u})$ of $\tilde{u}$ as

$$\begin{aligned}\Sigma_{\tilde{u}} =& GZ^\top CH_C^{-1}C\mathrm{cov}\big(y - X\hat{\beta}\big)CH_C^{-1}CZG\\ =& \sigma_\varepsilon^2 GZ^\top CH_C^{-1}CHCH_C^{-1}CZG\\ &+ \sigma_\varepsilon^2 GZ^\top CH_C^{-1}CX\big(X^\top CH_C^{-1}CX\big)^{-1}X^\top CH_C^{-1}CZG\\ &- 2\sigma_\varepsilon^2 GZ^\top CH_C^{-1}CHCH_C^{-1}CX\big(X^\top CH_C^{-1}CX\big)^{-1}X^\top CH_C^{-1}CZG\\ =& \sigma_\varepsilon^2 GZ^\top CH_C^{-1}CZG - \sigma_\varepsilon^2 GZ^\top CH_C^{-1}CX\big(X^\top CH_C^{-1}CX\big)^{-1}X^\top CH_C^{-1}CZG.\end{aligned}$$

The covariance matrix of $\tilde{u}$ is invariant to centring $X$ and $Z$. □

*Proof.* (Lemma 2) We use the unbiasedness of $\hat{\beta}$ and calculate the expectation of $\hat{R}_X^2$ with respect to $y$ and find

$$\begin{aligned}E(\hat{R}_X^2) &= E\big\{\hat{\beta}^\top\hat{\Sigma}_X\hat{\beta} - \mathrm{tr}\,\big(\hat{\Sigma}_X\Sigma_{\hat{\beta}}\big)\big\}\\ &= \mathrm{tr}\,\big\{\hat{\Sigma}_X E(\hat{\beta}\hat{\beta}^\top)\big\} - \mathrm{tr}\,\big(\hat{\Sigma}_X\Sigma_{\hat{\beta}}\big)\\ &= \beta^\top\hat{\Sigma}_X\beta,\end{aligned}$$

which implies that $\hat{R}_X^2$ is unbiased for the empirical variance $\beta^\top\hat{\Sigma}_X\beta$. Similar to Schreck et al. (2019), we calculate

$$E(\tilde{u}^\top\hat{\Sigma}_Z\tilde{u}) = E(\tilde{u})^\top\hat{\Sigma}_Z E(\tilde{u}) + \mathrm{tr}\,\big(\hat{\Sigma}_Z\Sigma_{\tilde{u}}\big) = \mathrm{tr}\,\big(\hat{\Sigma}_Z\Sigma_{\tilde{u}}\big).$$

We take the expectation of $\tilde{R}_Z^2$ with respect to $y$, and because $\hat{\Sigma}_Z$, $\Sigma_{\tilde{u}}$, and $D$ do not include $y$, we obtain

$$E(\tilde{R}_Z^2 - u^\top\hat{\Sigma}_Z u) = E(\tilde{u}^\top\hat{\Sigma}_Z\tilde{u}) - \mathrm{tr}\,\big(\hat{\Sigma}_Z\Sigma_{\tilde{u}}\big) + \mathrm{tr}(D\hat{\Sigma}_Z) - E(u^\top\hat{\Sigma}_Z u) = 0,$$

because $E(u^\top\hat{\Sigma}_Z u) = \mathrm{tr}(\hat{\Sigma}_Z D)$. This results implies unbiasedness for predictors, for detailed information we refer to Searle et al. (1992, §7), for instance. We use (15), (18) and (19) to calculate

$$\begin{aligned}\mathrm{cov}(\hat{\beta}, \tilde{u}) &= \mathrm{cov}\big\{\hat{\beta}, CH_C^{-1}C(y - X\hat{\beta})\big\}\\ &= \big\{\mathrm{cov}(\hat{\beta}, y) - \mathrm{cov}(\hat{\beta})X^\top\big\}CH_C^{-1}C\\ &= \big\{\big(X^\top CH_C^{-1}CX\big)^{-1}X^\top CH_C^{-1}CH\sigma_\varepsilon^2 - \big(X^\top CH_C^{-1}CX\big)^{-1}X\sigma_\varepsilon^2\big\}CH_C^{-1}C\\ &= 0,\end{aligned}$$

i.e. $\hat{\beta}$ and $\tilde{u}$ are uncorrelated. This leads us to

$$\begin{aligned}E\big(\hat{\beta}^\top\hat{\Sigma}_{XZ}\tilde{u}\big) &= \mathrm{tr}\,\big\{\hat{\Sigma}_{XZ}E(\tilde{u}\hat{\beta}^\top)\big\}\\ &= \mathrm{tr}\,\big\{\hat{\Sigma}_{XZ}E(\tilde{u})E(\hat{\beta}^\top)\big\} + \mathrm{tr}\,\big\{\hat{\Sigma}_{XZ}\mathrm{cov}(\tilde{u}, \hat{\beta})\big\}\\ &= \beta^\top\hat{\Sigma}_{XZ}E(\tilde{u}) = 0,\end{aligned}$$

which equals $E(\beta^\top\hat{\Sigma}_{XZ}u) = 0$. This proves the unbiasedness of the predictor $\hat{\beta}^\top\hat{\Sigma}_{XZ}\tilde{u}$. □

*Proof.* (Lemma 3.) We use (the dot variant of) Lemma 4 to write

$$\dot{\sigma}_{\varepsilon}^2 = \frac{1}{n-k-1} y^{\top} \dot{P}_H^I y = \frac{1}{n-k-1} y^{\top} C \dot{P}_H^C C y.$$

This implies that $\dot{\sigma}_{\varepsilon}^2$ is invariant to centring $y$, $X$ and $Z$ because $\dot{P}_H^C$ is independent of the centring of $X$ and $Z$. In addition, it holds that

$$\dot{P}_H^I y = \dot{H}^{-1}\big(I_n - \tilde{X}\big(\tilde{X}^{\top}\dot{H}^{-1}\tilde{X}\big)^{-1}\tilde{X}^{\top}\dot{H}^{-1}\big)y = \dot{H}^{-1}\big(y - 1_n\dot{\mu} - X\dot{\beta}\big). \tag{20}$$

We use that $\dot{H}\dot{P}_H^I$ is idempotent and write

$$y^{\top}\dot{P}_H^I y = y^{\top}\dot{H}^{-1}\dot{H}\dot{P}_H^I\dot{H}\dot{P}_H^I y = y^{\top}\dot{P}_H^I\dot{H}\dot{P}_H^I y.$$

Combining the last two results leads to

$$\dot{\sigma}_{\varepsilon}^2 = \frac{1}{n-k-1}\big(y - 1_n\dot{\mu} - X\dot{\beta}\big)^{\top}\dot{H}^{-1}\big(y - 1_n\dot{\mu} - X\dot{\beta}\big).$$

We use (17) to write

$$\begin{aligned}
\dot{\sigma}_{\varepsilon}^2 &= \frac{1}{n-k-1}\big(y - 1_n\dot{\mu} - X\dot{\beta}\big)^{\top}\dot{H}^{-1}\dot{H}\dot{H}^{-1}\big(y - 1_n\dot{\mu} - X\dot{\beta}\big) \\
&= \frac{1}{n-k-1}\big(y - X\dot{\beta}\big)^{\top}C\dot{H}_C^{-1}C\dot{H}C\dot{H}_C^{-1}C\big(y - X\dot{\beta}\big) \\
&= \frac{1}{n-k-1}\big(y - X\dot{\beta}\big)^{\top}C\dot{H}_C^{-1}C\big(y - X\dot{\beta}\big),
\end{aligned}$$

which holds because

$$C\dot{H}C = C + CZ\dot{D}Z^{\top}C = \dot{H}_C - \frac{1}{n}1_n1_n^{\top}$$

and

$$\dot{H}_C^{-1}1_n \overset{(5)}{=} 1_n - CZ\big(\dot{A} - n^{-1}Z^{\top}1_n1_n^{\top}Z\big)^{-1}Z^{\top}C1_n = 1_n.$$

The last equality results from $C1_n = 0$. We use Lemma 4 and equations (17) and (20) to obtain

$$\begin{aligned}
\dot{\sigma}_{\varepsilon}^2 &= \frac{1}{\operatorname{tr}\big(\dot{P}_H^I\big)} y^{\top}\dot{P}_H^I\dot{P}_H^I y \\
&= \frac{1}{\operatorname{tr}\big(C\dot{P}_H^C C\big)}\big(y - 1_n\dot{\mu} - X\dot{\beta}\big)^{\top}\dot{H}^{-1}\dot{H}^{-1}\big(y - 1_n\dot{\mu} - X\dot{\beta}\big) \\
&= \frac{1}{\operatorname{tr}\big(C\dot{P}_H^C C\big)}\big(y - X\dot{\beta}\big)^{\top}C\dot{H}_C^{-2}C\big(y - X\dot{\beta}\big).
\end{aligned}$$

Also, we find that

$$\begin{aligned}
&\dot{\sigma}_{\varepsilon}^2 \operatorname{tr}\big(\dot{P}_H^I Z_i Z_i^{\top}\big) = y^{\top}\dot{P}_H^I Z_i Z_i^{\top}\dot{P}_H^I y \\
\iff\ &\dot{\sigma}_{\varepsilon}^2 \operatorname{tr}\big(\dot{P}_H^C C Z_i Z_i^{\top} C\big) = y^{\top}C\dot{P}_H^C Z_i Z_i^{\top} C\dot{P}_H^I C y.
\end{aligned}$$

The restricted maximum likelihood estimators $\dot{\sigma}_{u_i}^2$ $(i = 1, \ldots, r)$, are invariant to centring $y$, $X$ and $Z$. □
